# Larval swimming in the sea anemone *Nematostella vectensis* is sensitive to a broad light spectrum and exhibits a wavelength-dependent behavioral switch

**DOI:** 10.1101/2023.11.10.566660

**Authors:** Meghan Payne, Emma Lilly, Camilla R Sharkey, Kyle J McCulloch

**Affiliations:** Department of Ecology, Evolution, and Behavior, University of Minnesota, St. Paul MN, 55108, U.S.A

**Keywords:** *Nematostella vectensis*, light behavior, opsin, Anthozoa, larval swimming, sea anemone

## Abstract

In nearly all animals, light sensing mediated by opsin visual pigments is important for survival and reproduction. Eyeless light-sensing systems, though vital for many animals, have received relatively less attention than forms with charismatic or complex eyes. Despite no single light sensing organ, the sea anemone *Nematostella vectensis*, has 29 opsin genes and multiple light-mediated behaviors throughout development and reproduction, suggesting a deceptively complex light-sensing system. To characterize one aspect of this light-sensing system, we analyzed larval swimming behavior at high wavelength resolution across the ultraviolet and visual spectrum. *N. vectensis* larvae respond to light at least from 315 to 650 nm, which is a broad sensitivity range even compared to many animals with complex eyes. Swimming in the water column is induced by ultraviolet (UV) and violet wavelengths until 420 nm. Between 420 and 430 nm a behavioral switch occurs where at wavelengths longer than 430 nm, larvae respond to light by swimming down. Swimming down toward the substrate is distinct from light avoidance, as animals do not exhibit positive or negative phototaxis at any wavelength tested. At wavelengths longer than 575 nm, animals in the water column take increasingly longer to respond and this behavior is more variable until 650 nm where larval response is no different from the dark, suggesting these longer wavelengths lie outside of their sensitivity range. Larval swimming is the only motile stage in the life history of *N. vectensis*, and increased swimming activity in the water column could lead to greater dispersal range in potentially damaging shallow environments with short-wavelength light exposure. Longer wavelength environments may indicate more suitable substrates for metamorphosis into the polyp stage, where the individual will remain for the rest of its life. Future work will test whether this robust behavior is mediated by multiple opsins.

## Introduction

Nearly all animals use visual systems to detect light signals from the environment, conspecifics, or from other species (Land and Nilsson, 2012; Terakita and Nagata, 2014). The evolution and function of complex eyes have historically received the most attention, with much of our knowledge of visual function and evolution coming from a few focal eye types (Arendt et al., 2009; Fernald, 2006; Nilsson, 2013). However species and life stages with simple or no eyes are common, and animals with complex eyes have multiple non-visual light-sensing systems (Kaniewska et al., 2015; Nordström et al., 2003; Porter, 2016). Yet we know much less about the molecular, cellular, and behavioral aspects of these light-sensing systems. Eyes likely evolved multiple times throughout evolution from more dispersed light sensing systems, and extant eyeless forms may hold important clues to understanding the repeated evolution of distinct eye types. To begin to address this major evolutionary question, we can ask how animals interface with the environment through behavior, as a first step in understanding light sensing traits in eyeless species.

Cnidaria (jellyfish, sea anemones, corals, etc.) are sister to Bilateria (vertebrates, arthropods, mollusks, etc.), making them an important phylogenetic comparison to better-known visual systems. The lineage contains a wide range of visual system complexity, from eyeless to camera-type eyes like our own. Medusozoa, the group including jellyfish and *Hydra,* have evolved eyes at least 8 times from eyeless forms (Picciani et al., 2018). Mostly correlational evidence suggests cnidarian eyes share some homology with bilaterian visual systems in their development, photoreceptors (opsins), and phototransduction pathways (Gehring, 2005; Hansen et al., 2023; Koyanagi et al., 2008; Kozmik et al., 2008, 2003). Some genetic links to light behavior in non-ocular light sensing have also been shown in Medusozoa. For instance, in the hydrozoan jellyfish *Clytia hemispherica,* a blue light absorbing opsin is responsible for oocyte maturation (Quiroga Artigas et al., 2018). *Hydra magnipapillata* are eyeless, yet evidence shows their contractile response and cnidocyte firing are regulated by light in a mechanism similar to some bilaterian phototransduction (Macias-Munõz et al., 2019; Plachetzki et al., 2012, 2010; Vöcking et al., 2022).

Anthozoa, the other major cnidarian group including corals, sea pens, and sea anemones, are all eyeless. Almost nothing is known about any molecular, genetic, or cellular processes of light sensing in this group, while behavioral data to specific light stimuli are similarly limited. Despite this lack of data, light-associated behaviors such as spawning are well-known in this clade (Boch et al., 2011; Fritzenwanker and Technau, 2002; Kaniewska et al., 2015; Lin et al., 2021). Much anthozoan behavior work has focused on circadian rhythms in feeding activity and spawning (Hendricks et al., 2012; Reitzel et al., 2013; Tarrant et al., 2019). Previous work has mostly tested behaviors in adults under natural light or dark conditions (Bell et al., 2006; Fabricius and Klumpp, 1995; Sebens and DeRiemer, 1977), but these do not attempt to parse salient light signals, and few have tested multiple signals or light conditions (Leach and Reitzel, 2020; Mason et al., 2011). Recently it has been shown that many species in the group including anemones and corals, the Hexacorallia, have very high numbers of opsins (McCulloch et al., 2023). As opsins are the protein component of photoreceptors in animal vision and in many forms of non-ocular light sensing, it is possible anemones and corals have unexplored complexity in their light-sensing systems. Thus, specific visual cues should be tested to better characterize the potentially complex light-mediated behavioral repertoire in Anthozoa at multiple life stages.

*Nematostella vectensis* is an emerging model sea anemone (Hexacorallia:Anthozoa) whose external fertilization, rapid development, robust whole-body regeneration, and high quality genomes make it amenable for laboratory study (Fletcher and Pereira da Conceicoa, 2023; Layden et al., 2016; Rentzsch et al., 2017; Zimmermann et al., 2020). *N. vectensis* is a shallow-water, estuarine, burrowing anemone, and can reproduce sexually or asexually via transverse fission (Layden et al., 2016). In sexual reproduction at room temperature, fertilized eggs begin to divide after about 2 hours, undergo gastrulation in under 24 hours, and develop into swimming larvae by about 48 hours. The swimming planula larva is the only dispersal stage of this otherwise sessile animal. After about a week the planula elongates, undergoes metamorphosis and forms a polyp, a miniature adult form with 4 tentacles surrounding a single opening (mouth). The polyp grows and adds up to 16 total tentacles and becomes sexually mature in three months (Layden et al., 2016). The adult polyp burrows into soft, shallow substrate in brackish water marshes and estuaries after metamorphosis where it will remain for the rest of its life.

Therefore, as the planula swims in the water column, finding a suitable substrate is crucial for an individual’s future survival and reproduction. To accomplish this task, the planula is uniformly covered in motile cilia that propel it through the water (Figure 1A), and it also has a sensory organ with much longer cilia called the apical organ located at the aboral (non-mouth pole) ectoderm oriented forward in the direction of swimming (Hand and Uhlinger, 1992; Marinković et al., 2020). This apical organ is required for metamorphosis, and expresses multiple homologous genes known in other species to be sensory-related, including opsins (McCulloch et al., 2023; Rentzsch et al., 2008; Sinigaglia et al., 2015). Despite no eye, *N. vectensis* is typical among hexacorals in having among the most opsins of any animal, at 29 (McCulloch et al., 2023). The expression patterns of these opsins and presence of genes involved in multiple bilaterian phototransduction pathways suggest a diversity of light-mediated behavioral responses in *N. vectensis* (Hansen et al., 2023; McCulloch et al., 2023). For instance, some opsins are found only in adults and may be sexually dimorphic in expression levels, suggesting a role in reproduction, while others are only found in larval stages, suggesting a role in swimming and substrate finding (McCulloch et al., 2023). However nothing is known about the sensory mechanisms required for substrate finding.

**Figure 1.**
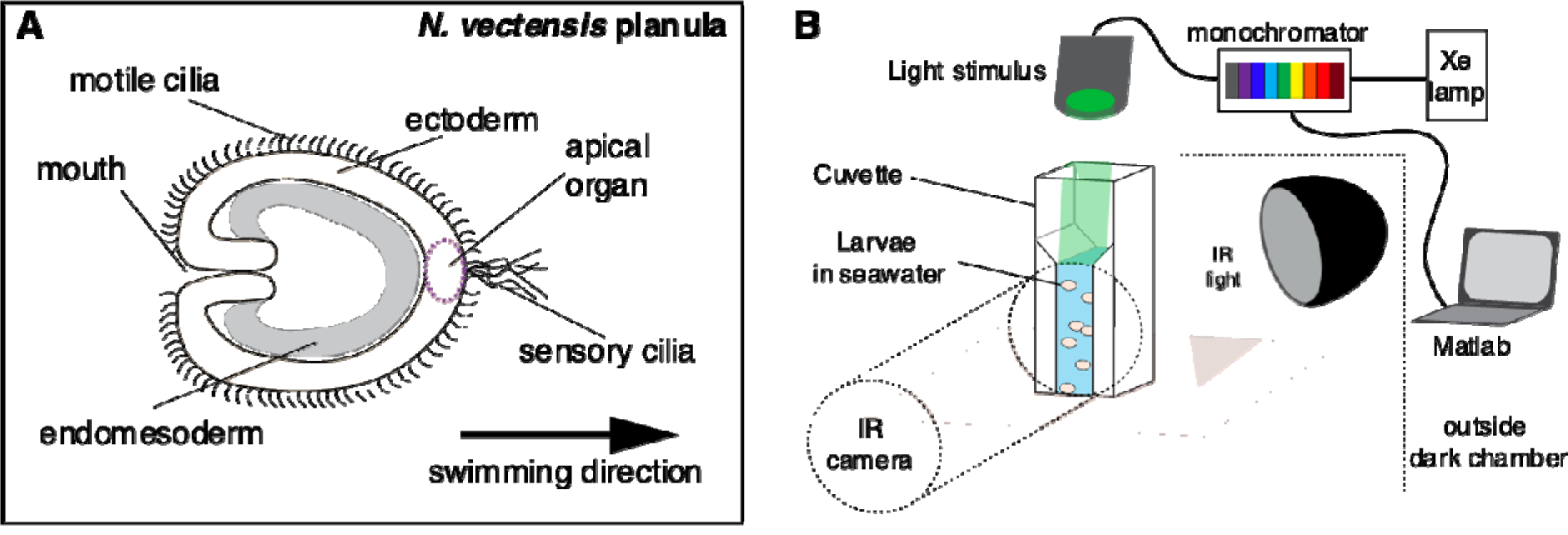
Larval anatomy and behavioral setup. **A)** Planulae are diploblastic and covered uniformly in motile cilia. Swimming direction is opposite to the oral pole, with a sensory ciliated organ known as the apical organ at the front in the direction of swimming. **B)** Experimental setup. Light from a xenon arc lamp passes through a monochromator controlled by a computer and is delivered via fiber optic cable into the dark chamber directly above the cuvette. The animals are bathed in infrared light and the infrared camera faces the cuvette from the side.

To begin to understand the connections between opsins and light sensing while swimming, we need a detailed understanding of the behavioral phenotype. Larval swimming in *N. vectensis* is known to increase with “disturbance” including to light, but this behavior is uncharacterized in the literature (Hand and Uhlinger, 1992). To characterize this light-mediated behavior, we tested the response of *N. vectensis* larvae to 54 monochromatic wavelengths of light in *N. vectensis* larval swimming. We show *N. vectensis* larvae can detect wavelengths at least between 315 and 650 nm, and larvae respond to these colors (UV to red) in consistent and robust ways.

## Materials and Methods

### Husbandry

*Nematostella vectensis* adults were kept in ⅓ concentration Instant Ocean seawater, at a constant temperature of 18℃, alternating light and dark exposure every 12 hours. Adults were fed brine shrimp 4-5 times a week. To induce spawning, *Nematostella vectensis* were placed in containers exposed to 24℃ and white light for 6 hours overnight. Eggs were collected in the morning, fertilized, and allowed to develop for 48 hours at room temperature. At two days post-fertilization larvae were placed in a petri dish in the dark at 18℃ for at least 24 hours before the experiment.

### Short wavelength Behavior Experimental Setup (315 - 550 nm)

Behavior experiments were conducted in a dark chamber with blackout curtains on an air table. See Figure 1 for behavior setup schematic (Figure 1A). Hardware for generation of light stimulus were located outside of the chamber and stimulus was delivered via fiber optic cable. The intensity and wavelength of light from a 150W xenon arc lamp (Osram) was controlled using a monochromator (Cairn Research) and a DAQ board (National Instruments), using custom MATLAB scripts (Data Acquisition Toolbox, Mathworks). Wavelength was determined by the angle of a 2400 line-ruled diffraction grating (315 – 550 nm) or 1200 line-ruled grating (450 – 700 nm) and intensity was adjusted by changing the width of monochromator in and out slits. The output light was measured at the approximate point of animal illumination using a spectrophotometer (Ocean FX, OceanOptics) calibrated for irradiance measurements (DH-3P-BAL-CAL, OceanOptics). Photon flux and wavelength were measured and adjusted to yield isoquant stimuli (7.18 × 10^15^ photons cm^−2^ s^−1^), at each test wavelength. See Sharkey *et al*. (2020) for further information about the optical set up and monochromator calibration.

Between 100 - 200 dark-adapted larvae were added with sea water in a cuvette with the light source directly above. Animals were allowed to settle in the cuvette in the dark for 30 minutes prior to the start of the experiment. The stage was lit for filming with an A14 infrared illuminator peaking at 850 nm (Tendelux). Dim red light was used to work in the dark room. Light was delivered automatically via Matlab using the Data Acquisition Toolbox (Mathworks), maintaining intensity (photon flux) while changing wavelength. Experiments either alternated between 10 minutes of dark and 10 minutes of light periods, or continuous wavelength changes with no dark periods. Light behavior was indistinguishable between the two methods, so we combined averages from both for our short wavelength dataset. The long wavelength dataset alternated between UV and light or dark treatment in randomized order. For complete experimental information all of our light protocols and raw tracked data are available on Dryad: https://doi.org/10.5061/dryad.wdbrv15vs. The order of wavelengths was semi-randomized to ensure both short and long wavelengths for normalization (see below). The same batch of animals was not re-used for any single wavelength, though the same batch could be used again to be able to record all wavelengths over two or three days of experiments. The cuvette was filmed from the side with a Minolta MN200 NV infrared video camera, in the dark, with infrared light for illumination at 30 frames per second.

Pilot experiments were carried out before the dataset was collected, and temperature sensors were used to record changes in temperature over the course of the different wavelengths. The probe was placed in the seawater in the cuvette in identical conditions to our experimental dataset. No changes were noted in temperature at any wavelength measured to at least one hundredth of a degree Celsius.

### Long Wavelength Behavior Experimental Setup (575 - 700 nm)

Longer wavelength experiments were conducted using a similar setup as above with the following changes. The monochromator used to generate longer wavelengths was unable to also generate short-wavelength light needed for the larvae to swim into the water column (see Results). To be sure that larvae were responding to longer wavelengths, we used an LED UV light controlled by a dual OptoLED power supply (Cairn Research) which peaked at 370 nm. The output of the LED was measured using a spectrophotometer at the point of the animal and manually adjusted until it was set to the same approximate total intensity as the long wavelengths presented to the animals. Animals were exposed to 5 minutes of UV light, then the test wavelength (575, 600, 620, 675 or 700 nm), or darkness for one hour. We saw no difference in swimming behavior in preliminary tests with one or two hours in the dark, therefore only one hour of darkness was used for experiments. For wavelengths 575 nm and 600 nm, behavioral response reached equilibrium (all animals swam to the bottom of the cuvette) well before an hour and some trials were stopped earlier than an hour.

### Video Analysis

Video analysis of the swimming larvae was conducted using FIJI v2.9.0/1.53t and Trackmate v7.0 (Ershov et al., 2022; Schindelin et al., 2012). Examples of input video screenshots are shown in Figure 1B. Videos were imported by the ffmpeg plugin v4.2.2 in FIJI and decimated by 100 to reduce file size for analysis. The region of the cuvette to be analyzed was defined using the FIJI selection tool, avoiding the walls of the cuvette, the water surface, or the bottom where larvae are settled (see Figure 1B). Trackmate was run with default settings except for the quality threshold, which was changed to 2.4. Trackmate calculated the number of spots within the region of interest corresponding to each swimming larva anywhere in the water column, at every frame. The number of spots per frame was exported as a .csv file and then normalized by the peak spot number over the entire experiment to get normalized swimming activity (i.e. 1 would indicate the most larvae swimming in the water column and 0 would indicate all larvae settled on the bottom of the cuvette). At all wavelengths, the longest it took to adjust from the previous condition and reach the new wavelength equilibrium was about 90 analysis frames (5 minutes, examples Figure 3). Therefore, averages were calculated after equilibrium was reached (after 90 frames). Averages from each replicate experiment and wavelength were then averaged and plotted as a box and whisker plot between 315 nm and 550 nm (See Appendix 1 for replicate numbers). After data collection during analysis, some data were removed if identical conditions were not met over the course of the entire video, (e.g., cuvette was moved during trial, not positioned directly facing camera, too few or too many animals, etc.). To quantify significant differences in this large dataset, the data were tested for normality using a Shapiro-Wilk test (W = 0.93911, p-value < .00000001), followed by a Kruskal-Wallis rank sum test to determine if there was a significant effect of wavelength on the response (χ^2^ = 228.58, d.f. = 47, p-value < 1 × 10^15^). To determine which treatments were significantly different from one another a pairwise Wilcoxon rank sum exact test was performed with Benjamini-Hochberg (BH) correction for multiple tests (Table 1).

To show that the response of larvae to darkness was not the same as lack of short wavelengths, we binned examples from our entire dataset where wavelengths went from short to long (N = 9), or long to short (N = 7), with dark periods in between. “Short” was defined as any wavelength shorter than 420 nm, and “long” was defined as any wavelength longer than 430 nm. The average swimming activity from the dark periods immediately preceding and following were compared with the activity for either the long or short wavelength in between. A Shapiro-Wilk test showed both short-to-long (W = 0.81547, p-value < 0.00001) and long-to-short (W = 0.81731, p-value = 0.001) datasets were non-normally distributed, and Kruskal-Wallis tests showed both had significant differences in the data (short-to-long, χ^2^ = 17.429, d.f. = 2, p-value < 0.001; long-to-short, χ^2^ = 13.781, d.f. = 2, p-value = 0.001). Wilcoxon rank sum exact tests were performed with Benjamini-Hochberg correction for multiple tests for each condition (Tables 2,3).

To calculate timing to equilibrium, swimming activity over time for each trial was fitted with a sigmoid curve using non-linear least squares (nls) regression, using the function:

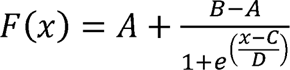

where the minimum asymptote (A), the maximum asymptote (B), the x-value at the midpoint between A and B (C), and the slope value (D) were all predicted given the data. We used the value for C to estimate halfway to settling time and plotted these values as box plots. At wavelengths greater than 650 nm, behavior was variable and often unresponsive, so sigmoid fits were not appropriate. We compared half-times of the swimming downwards behavior at longer wavelengths (575, 600, 625, 650 nm) with representative shorter wavelength downward swimming responses estimated by the same method (450 nm, 500 nm). The Shapiro-Wilk test for normality (W = 0.77472, p-value < 0.0001) showed the data were not normally distributed. A Kruskal-Wallis rank sum test was performed (χ^2^ = 21.557, d.f. = 5, p-value < 0.001), followed by a pairwise Wilcoxon rank sum exact test was performed with BH correction to determine which treatments were significantly different from one another (Table 4).

To quantify lack of responses at longer wavelengths, for each trial, the last 20 frames were averaged and subtracted from the average of the first 20 frames at each wavelength. This yielded the change in swimming activity after one hour of light exposure. These data were plotted as box plots and compared similarly to above. The Shapiro-Wilk test for normality (W = 0.89426, p-value < 0.001) showed the data were not normally distributed. A Kruskal-Wallis rank sum test was used to determine if there were significant differences among treatments in the data, which there were (χ^2^ = 14.955, d.f. = 6, p-value < 0.05). To determine which treatments were significantly different from one another a pairwise Wilcoxon rank sum test was performed with BH correction for multiple tests (Table 5).

### Phototaxis Experiments

Phototaxis experiments were conducted as above, but with the cuvette turned on its side. The light stimulus was applied at one end of the cuvette (Figure 2D). Larvae were allowed to spread across the full length of the cuvette in the dark for 30 minutes. After 30 minutes, the light protocol was started, and larval behavior recorded in infrared. Video analysis was similar to above, but instead of total number of spots, the average × position of total spots for any given wavelength was used from the Trackmate output data. No changes in average × position were seen at any wavelength tested.

**Figure 2.**
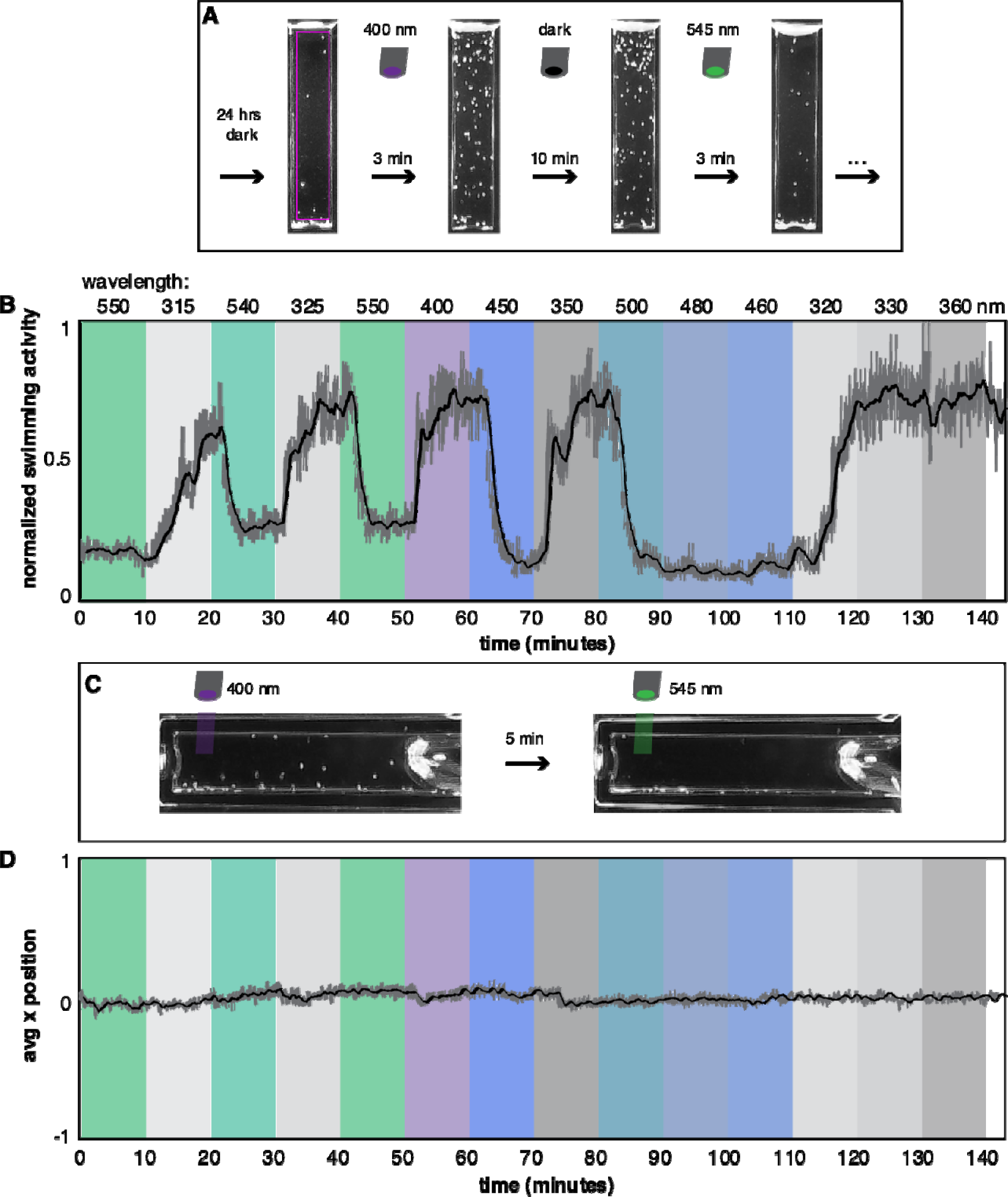
Experimental setup and example video analysis. **A)** Example video frames analyzed by Trackmate, showing differences between light conditions. After being in the dark for 24 hours, animals are not swimming in the water column. In UV light, swimming immediately increases within 3 minutes. Swimming activity is not changed in the dark following UV light after 10 minutes (or at least 2 hours, see below). Green light causes nearly all larvae to swim down within 3 minutes. Pink box shows region of interest where swimming activity is tracked. **B)** Example plotted after Trackmate analysis with wavelength treatments overlaid on the experiment in color. Downward swimming occurs with any wavelength over 430, swimming into the water column occurs at any wavelength shorter than 410. Dark gray line is raw data, black line is moving average over 20 previous datapoints. **C)** Example of phototaxis setup under two light conditions. Animals do not swim on average toward or away from light source, but still show up and down swimming in the water column. **D)** Example results from a single phototaxis experiment. Total of 7 experiments were conducted. Average x position of the spots calculated from Trackmate does not change and remains evenly distributed across x positions (near 0), regardless of wavelength.

## Results

We conducted a multi spectral behavioral experiment to test the effect of light on the swimming behavour of larval *N. vectensis* (Figure 1B). Screenshots of typical videos analyzed are shown in Figure 2A. We define swimming activity as the normalized number of animals in the water column (within the pink box, Figure 2A). When presented with light between 315 nm and 420 nm, animals actively swam into the water column the entire light demonstration (Figure 2). This movement was not directionally toward the light or surface, but animals consistently moved in all directions throughout the water column. When animals were presented with light from ∼430 nm to 625 nm, they actively swam down to the bottom of the cuvette (as opposed to passive sinking) and continued swimming at the bottom. In contrast, after 24 hours of darkness, animals were not seen in the water column and were less active in general. We could not quantify differences in swimming activity at the bottom of the cuvette with our setup.

To further characterize and formally quantify this behavior, we measured photo-activity at 5 nm intervals under illumination from 315 to 550 nm. We first noticed visually obvious changes in swimming activity between short (< 425 nm) and longer (> 425 nm) wavelengths (Figure 2A,B). The behavioral response to either short or long wavelength was robust regardless of light/dark combinations, order of wavelengths, and up to at least five hours of testing (Figure 2B). Animals in the region of interest in the water column (pink box, Figure 2A) were tracked and defined as swimming “up” (Figure 2B). When animals moved to the bottom of the cuvette, they exited the region of interest, and these were labeled as swimming “down.”

We additionally wanted to know whether animals were moving toward or away from light (phototaxis). To assess whether swimming behavior was phototactic, the cuvette was placed on its side with the light on at one end (Figure 2C). Over a total of 8 experiments, animals did not move toward or away from the light in any wavelength tested, suggesting no phototaxis. Photo-activity (i.e. active swimming in the water column) was still present, similar to when the cuvette was upright only (Figure 2C,D).

To calculate levels of swimming activity under each light condition, we allowed larvae to achieve equilibrium, i.e. when the change in number of tracked larvae over time plateaued. This allowed us to empirically determine that larvae take less than two minutes on average to adjust to a new light condition when swimming up, while swimming down takes longer, at just under four minutes (Figure 3). Animals responded by swimming up in wavelengths as short as 315 nm (limit of monochromator) (Figure 4). At 335 nm through 415 nm, swimming activity was high (i.e. the majority of animals were “up,” or swimming in the water column), and not significantly different from one wavelength to another (Table 1). Interestingly, swimming activity at 335 nm is significantly higher than most other wavelengths, and wavelengths 315-325 nm are significantly lower than some other short wavelength activity levels (Figure 4, Table 1). At about 415 nm, swimming activity begins to decrease until about 425 nm, where a steep drop in activity occurs. At 425 nm there is a switch in behavior where larvae respond by actively swimming down (Figure 4). Between 415 nm and 425 nm, patchy significant differences with both short and long wavelengths suggests a transition zone between up and down swimming under blue-violet light (Figure 4, Table 1). Levels of swimming down are not significantly different from each other between 430 nm and 550 nm (Figure 2, Table 1). Lower swimming activities are not zero because at any given time, individuals may “jump” up into the water column and are counted, but then quickly descend back to the bottom.

**Figure 3.**
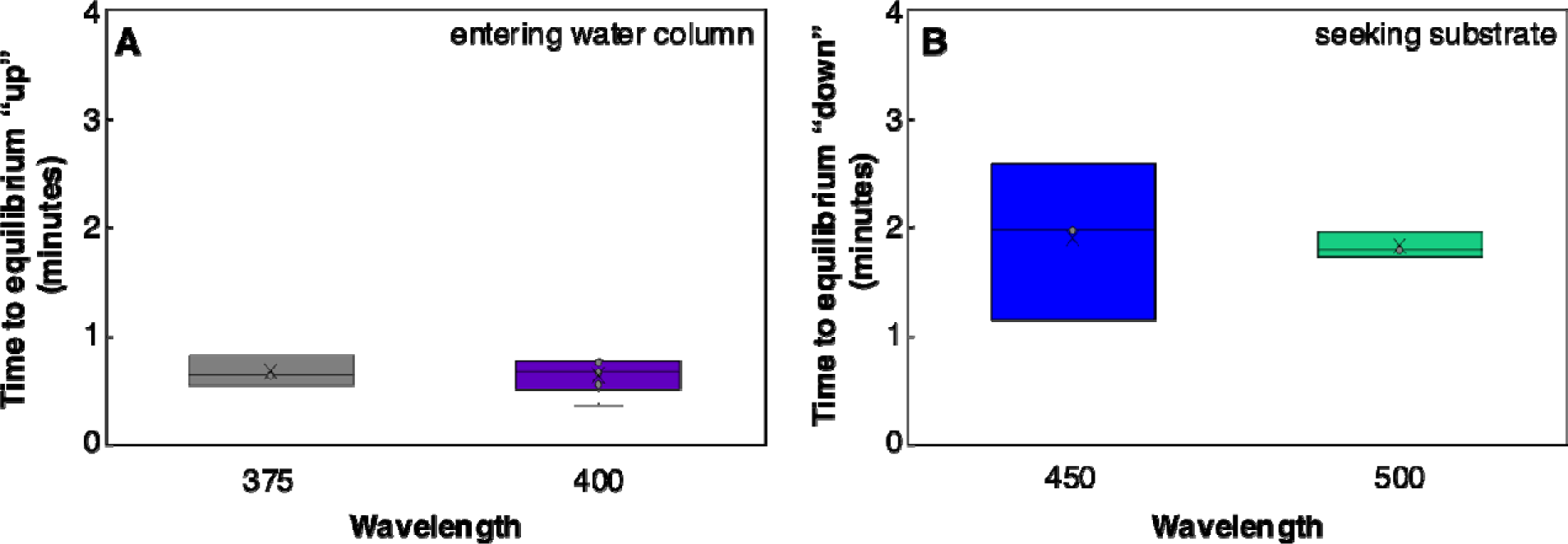
Timing of behavioral changes. Half time to behavioral shift from down to up (A) or up to down (B) is shown, as estimated by sigmoid curve fits for representative wavelengths. **A)** shows half time to peak swimming activity for wavelengths shorter than 425 nm. (375 nm, N = 4; 400 nm, N = 6). **B)** shows half time to peak down swimming activity for wavelengths longer than 425 nm (450 nm, N = 3; 500 nm, N = 3).

**Figure 4.**
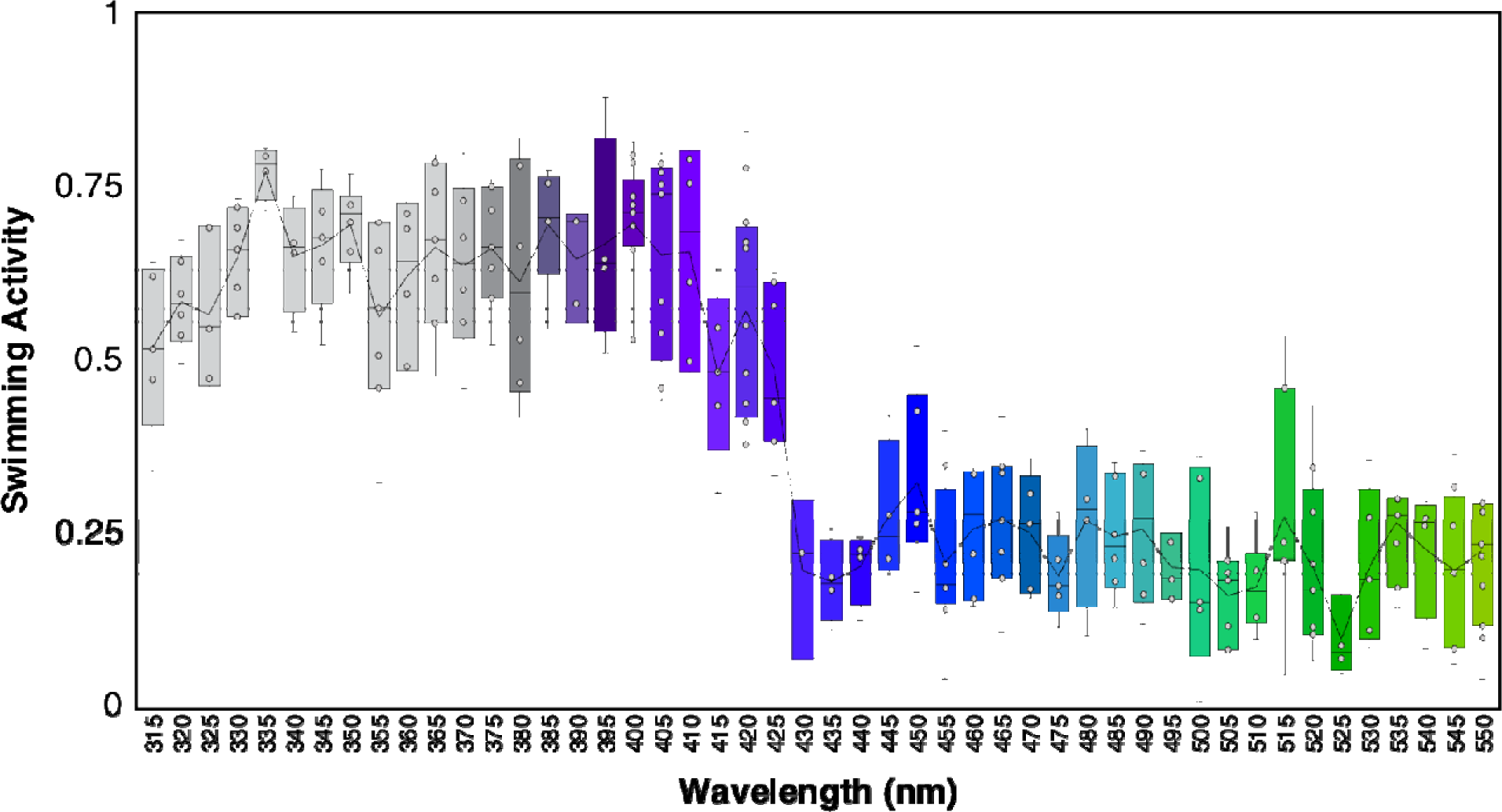
Average swimming activity from 315-550 nm. Boxplot of median normalized light behavior swimming activity levels by wavelength. Line connecting each wavelength represents the mean. Swimming activity at each wavelength was averaged over at minimum three experiments and up to seven experiments. A large decrease in swimming activity is seen between wavelengths 420 and 430. Number of experiments for each trial are shown in Appendix 1.

It is possible animals were responding to only a subset of wavelengths and the opposite behavior could be due to an absence of the salient signal. To show that animals were in fact responding to the broad range of wavelengths tested, we compared dark levels of swimming activity before and after light treatments. To show this, we binned all examples from our dataset where the sequence of stimuli tested went from short wavelength (315-415 nm) to dark, to long wavelength (430-550 nm), to dark again, or vice versa (long to dark to short to dark). The initial light stimulus would prime the animals to be maximally either up or down, and then the next stimulus should cause the opposite behavior, providing for the greatest change in swimming activity between any two stimuli. If downward swimming behavior in long wavelengths was similar to that in darkness, then we should see no difference in dark periods before or after long wavelength stimuli. Similarly, if the absence of light was causing animals to swim downwards then there should be a significant difference between dark periods before and after short wavelength stimuli. We show in both down-to-up and up-to-down sequences, swimming activity in the dark does not significantly change after any light stimulus (Wilcoxon rank sum test, p-value = 1; Tables 2,3), while dark swimming activity preceding the light stimulus is significantly different for short (Wilcoxon rank sum test, p < 0.001; Figure 5, Table 2) and long wavelengths (Wilcoxon rank sum test, p < 0.01; Figure 5, Tables 2,3). This shows that swimming behavior in the dark is dictated by the previous wavelength tested, and therefore animals sense and alter their behavior in response to both short and long wavelengths.

**Figure 5.**
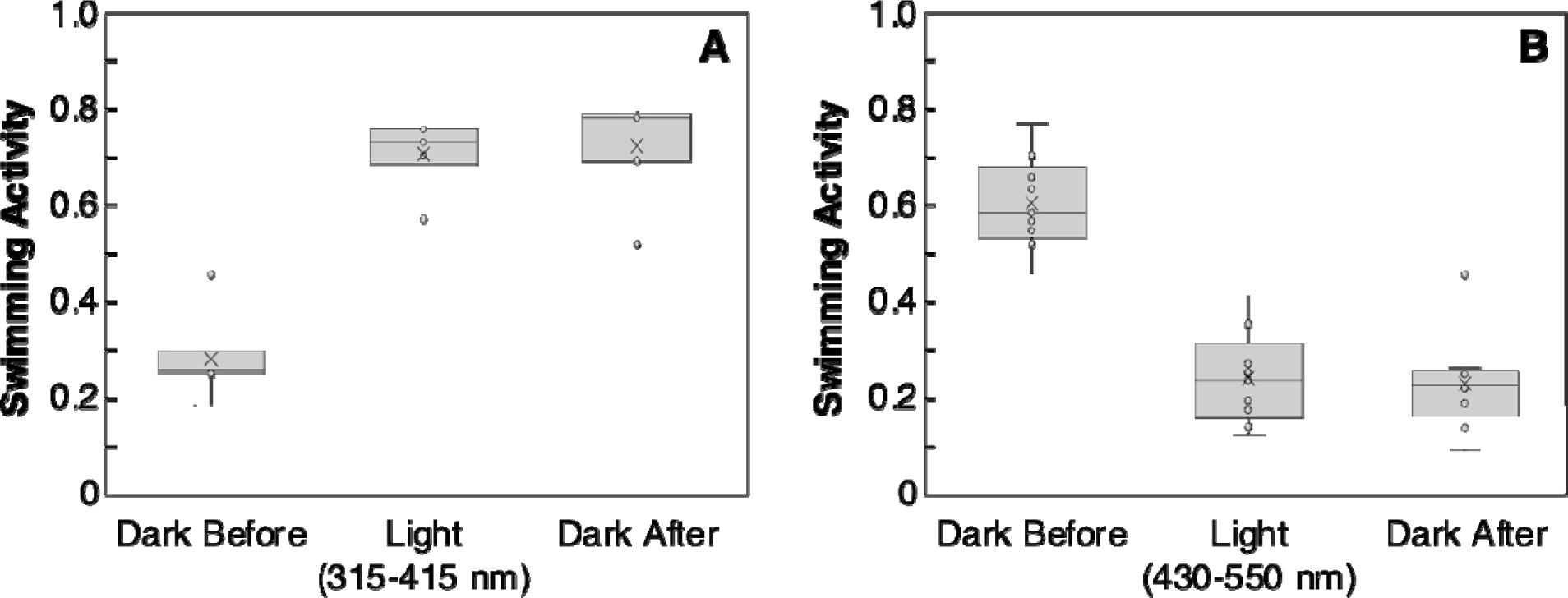
Short-term dark behavior is dictated by previous light stimulus. **A)** Swimming activity is shown for 10-minute periods under darkness followed by short wavelength light (315 – 415 nm), followed by darkness. The dark period before the stimulus is significantly different from either the light period or the dark period after the light (Wilcoxon rank sum test, p-value < 0.001). All trials with the sequence: long, dark, short, dark were binned and plotted (N = 7 experiments, first light stimulus not shown). **B)** Swimming activity is shown for darkness before the light stimulus, then long wavelength light (430 – 550 nm) then darkness. Darkness before the light stimulus is significantly higher than either the light stimulus or dark after (Wilcoxon rank sum test, p-value < 0.01). All trials with the sequence: short, dark, long, dark were binned and plotted (N = 9). See methods and Tables 2 and 3 for statistical tests and p-value tables.

For longer wavelengths (575 - 700 nm), changes in behavioral responses and technical limitations required us to modify the experiment (see Methods). We first noticed response time increased with longer wavelengths making it a challenge to analyze the data in ten-minute trials. We therefore increased trial time for these wavelengths, and to visually compare the change in responses with longer wavelengths we show the average entire activity over time at each wavelength (Figure 6). Downward swimming behavior at 575 nm begins immediately and has the steepest slope of the longer wavelengths, while at 600 nm, animals take longer to respond, and the slope is shallower (Figure 6A-C). At 625 nm, animals take even longer to respond, but still settle within one hour (Figure 6D). At 650, 675, and 700 nm, animals did not settle for at least an hour, although the slopes were trending downward (Figure 6E-G). In complete darkness, no change was noted after an hour (Figure 6H).

**Figure 6.**
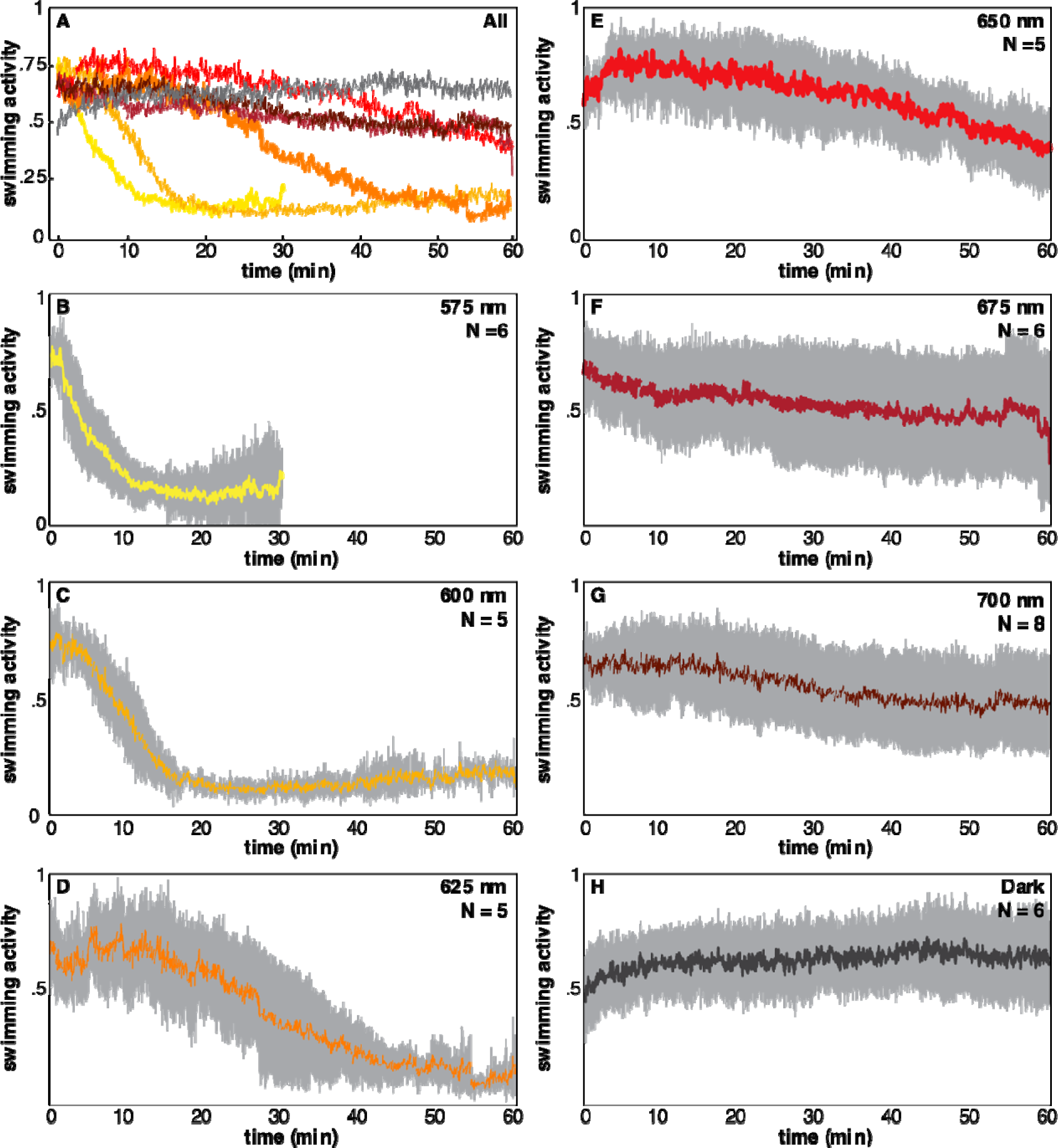
Swimming down response slows at longer wavelengths. **A)** All averaged responses over time for long wavelengths overlaid for comparison. Colors of individual lines correspond to colors and wavelengths in B-H. Averages are taken for at least 5 experiments per wavelength. **B-H)** The same data as in A, but data for each wavelength are shown individually with 95% confidence intervals. Each wavelength is labeled for each graph and colors correspond to colors in A.

We wanted to quantify these differences to better characterize the behavioral response. To do so, we fit data from each experiment with a sigmoidal function, identified the inflection point (half-time to down equilibrium) (See Appendix 2 for visualizations of curve fits). We then compared each long wavelength time with each other and two wavelengths from the short wavelength dataset (Figure 7A). We found that timing to down equilibrium is not significantly different between 450 nm, 500 nm, and 575 nm (Table 4). There is a significant increase in time from 575 to 625 nm, while 625 and 650 nm are not significantly different (Table 4). Despite no significance, 650 nm appears to continue to increase in down response time, but the variance in the data also increases with increasing wavelength (Figure 7A). Without reliable responses, we were unable to fit sigmoid curves to wavelengths longer than 650 or in the dark. To try to quantify differences in responses across all long wavelengths we instead took the difference at the beginning of the hour trial period and the end of the hour for each wavelength (Figure 7B). The trend of the data shows decreased difference between start and end time as well as increased variability in the data at longer wavelengths, with dark activity being the most variable and the median of the data showing a slight *increase* in activity at the end of the trial (Figure 7B). Despite these trends, the only significant difference in these data is between 575 nm and dark trials (Table 5).

**Figure 7.**
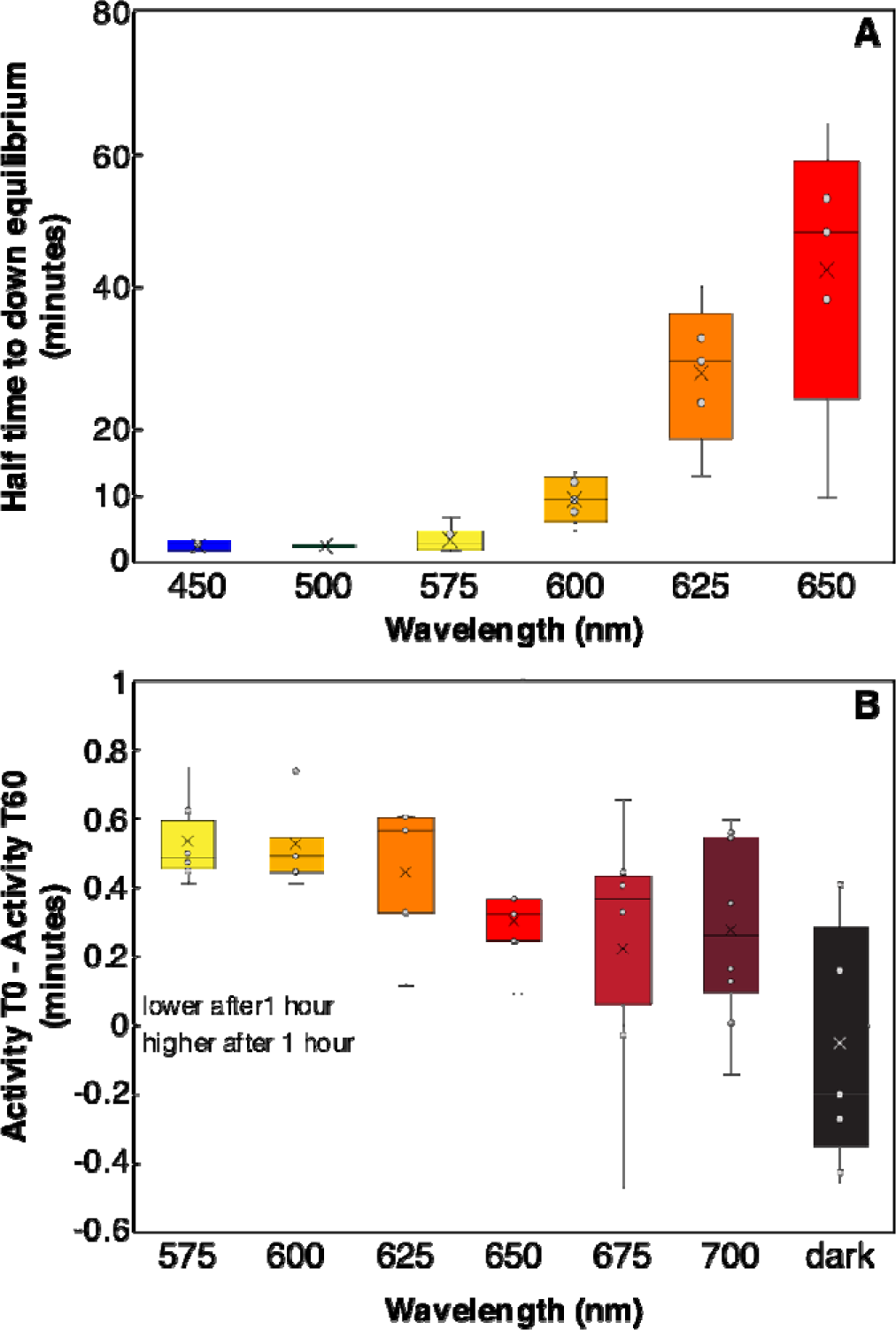
Swimming down response slows and resembles dark behavior at longe wavelengths. **A)** Half time to down equilibrium is based on estimates from sigmoid curve predictions (See methods, Appendix 2). At each of the longer wavelengths measured after 575 nm, settling response time tends to increase and becomes more variable with increasing wavelength. For significance values see Table 4. **B)** Data longer than 650 nm and dark data could not be fit with sigmoid curves so data were analyzed by measuring the difference in initial swimming activity and activity after 60 minutes. More positive values mean larvae swam down more, while negative values mean more larvae were in the water column at the end of the trial than at the beginning. A zero value would mean no change in between initial and final swimming activities. For significance values see Table 5.

## Discussion

Our results indicate that swimming larvae of *N. vectensis* can detect and respond to a broad range of wavelengths, from at least 315 nm to 625 nm. Larvae do not exhibit phototaxis, as they neither swam toward or away from the light in any wavelength tested. Instead, larvae switch between two swimming behaviors “swimming in the water column” and “swimming down toward substrate” at different wavelengths but do not swim directionally toward the surface or toward light. Swimming activity is high in wavelengths from 315 nm to 400 nm, but between 415 nm and 430 nm, high swimming activity transitions toward downward swimming. Animals respond to wavelengths from 430 nm up to at least 625 nm by swimming down toward the substrate. At wavelengths from 650 to 700 nm animals may have some weak response to light stimuli but the quantified behavior is indistinguishable from dark trials, suggesting this is the limit of larval sensitivity. Despite no eye, pigment, or complex sensory organ, the range of wavelength sensitivity of *N. vectensis* larvae is broader than the range of human color vision range, by comparison.

The range of wavelengths sensed by *N. vectensis* larvae suggests, assuming opsin-based light detection, that multiple opsins must be involved in this behavior. A single visual pigment would not be able to facilitate two distinct swimming behaviors (swimming in the water column and downward swimming). We have shown that neither behavior is simply induced by a lack of sensitivity to short or long wavelength light. Rather, animals exposed to darkness or wavelengths above 625 nm maintain their swimming behavior. Based on known opsin absorbance spectra (Stavenga, 2010), at least three opsins would be needed to cover this broad sensitivity range. This could be accomplished by a short-wavelength opsin or two, which when absorbing light, induce swimming in the water column via directional ciliary beating. Long wavelength opsins may induce a switch in cilia beating direction resulting in swimming down toward the substrate. A transition where two opsins may be sending conflicting signals could be between 415 and 430 nm, where some combination of larvae swim up and some swim down. We identified equilibria at these wavelengths that were neither fully active nor fully down, and there could be a choice individuals make to swim up or down, and this causes the population level average to be intermediate. It is unclear whether this response is specific to each individual and is fixed, or if an individual has some rate of changing its response, leading to this intermediate measured activity. Future experiments could test with fewer individuals to be able to track them individually. Future directions will address which opsins are involved through targeted CRISPR/Cas9 knockouts and behavioral assays like those established in this study. From previous work, one opsin, *NvASOII-8b,* is shown to be highly expressed in the sensory apical organ at larval stages (McCulloch et al., 2023). This could be responsible for part of the wavelength-specific response.

At wavelengths longer than 550 nm, the length of time it takes to respond to particular wavelengths increases. This could be that there is a long-wavelength absorbing rhodopsin whose peak absorbance is in the green range, and it is less able to absorb photons at these long red and near-infrared wavelengths. Thus, despite the same total intensity of stimulus, longer wavelengths are in effect dimmer and the behavioral response is therefore less sensitive. Although, not significantly different, the variances of the responses at 650 nm, 675 nm, and 700 nm, do appear different from complete darkness, suggesting there is potentially some sensitivity even to 700 nm light. At all longer wavelengths (625 - 700 nm), and in the dark, there were at least one or two experiments where animals settled quickly. In the dark, we know animals eventually will settle over a 24-hour period, and we saw some settle in the dark much before that, so there could be some other sensory cue or lack thereof that induces settling.

The sensitivity and robustness of this behavioral switch across such a broad range of wavelengths is unexpected for a “simple” eyeless and unpigmented larva, and it is unknown why this sensitivity would be required. One explanation is that these are shallow water species and are therefore subject to the full range of wavelengths from the sun (Pope and Fry, 1997; Smith and Baker, 1981). The larval stage is the only dispersal stage in the lifecycle of *N. vectensis* before metamorphosing into a sessile polyp. It could be that short wavelength light in shallow water is itself damaging to the polyp, or is an indicator of a harsh intertidal zone that would dry out polyps that are exposed at low tide. Without pigment or other means for directional swimming, simply swimming more would result in increased dispersal, on average, and could result in larvae reaching more favorable light environments for seeking substrate. In coastal shallow nutrient-rich waters, UV light is almost completely attenuated within a meter or two below the surface, and longer red wavelengths are attenuated within 5 or 10 meters, while blue-green light penetrates much further (Levine and MacNichol, 1982). In the presence of blue, green, or even red wavelengths and the absence of UV/violet, it may be beneficial to swim down and find suitable substrate on which to undergo metamorphosis.

Although the visual systems of eyeless animals have received less attention than their eyed relatives, our work shows that light sensing is important and can still be part of a complex trait by these eyeless species. We note that detailed characterization of sensory systems and species previously regarded as “simple”, such as anthozoan cnidarians, can reveal overlooked complexity in sensory traits and behavior. Further work will investigate the molecular-genetic basis for this and other behavioral traits to get a fuller comparative picture of visual system evolution. Although more work needs to be done, elucidating part of the *N. vectensis* visual system is an important step in broadening our understanding of the evolution of complex traits like animal eyes.

## Data Accessibility Statement

Trackmate raw data for all swimming activity used in this study is available in Dryad at https://doi.org/10.5061/dryad.wdbrv15vs. Videos are available upon request. All other data in this study are included in the figures or as appendices.

## Author Contributions section

KJM, conception and experimental design, acquisition of data, analysis, and interpretation of data, writing and editing manuscript. MP, experimental design, acquisition of data, analysis, and interpretation of data, writing manuscript. EL, experimental design, acquisition of data, analysis of data, editing manuscript. CRS, experimental design, writing and editing manuscript.

## Acknowledgements, including details of funding bodies with grant numbers

We’d like to thank Trevor Wardill for graciously allowing use of equipment, Paloma Gonzalez-Bellido and the Wardill and Fly-Sy labs for critical advice and input. Thanks also to Emilie Snell-Rood, Tim Mitchell and Luis Santiago-Rosario and the Snell-Rood lab for discussions about the manuscript. This work was supported by the University of Minnesota School of Biological Sciences, the University of Minnesota Office for the Vice President of Research (Grant in Aid of Research, New Faculty), by the University of Minnesota’s Office of Undergraduate Research (UROP), and by the UMN College of Biological Sciences’ Dean’s Research Program.

