## Appendices for "Larval swimming in the sea anemone *Nematostella vectensis* is sensitive to a broad light spectrum and exhibits a wavelength-dependent behavioral switch"

**Appendix 1: Number of replicates for short and long wavlength experiments**

| Wavelength | N experiments |
|---|---|
| 315 | 5 |
| 320 | 6 |
| 325 | 6 |
| 330 | 7 |
| 335 | 4 |
| 340 | 4 |
| 345 | 5 |
| 350 | 6 |
| 355 | 7 |
| 360 | 6 |
| 365 | 7 |
| 370 | 6 |
| 375 | 7 |
| 380 | 6 |
| 385 | 5 |
| 390 | 5 |
| 395 | 4 |
| 400 | 13 |
| 405 | 9 |
| 410 | 6 |
| 415 | 5 |
| 420 | 12 |
| 425 | 7 |
| 430 | 3 |
| 435 | 4 |
| 440 | 4 |
| 445 | 4 |
| 450 | 6 |
| 455 | 8 |
| 460 | 6 |
| 465 | 7 |
| 470 | 5 |
| 475 | 5 |
| 480 | 4 |
| 485 | 6 |
| 490 | 6 |
| 495 | 6 |
| 500 | 5 |
| 505 | 7 |
| 510 | 6 |
| 515 | 7 |

| | | |
|---|---|---|
| | 520 | 9 |
| | 525 | 4 |
| | 530 | 5 |
| | 535 | 7 |
| | 540 | 4 |
| | 545 | 8 |
| | 550 | 11 |
| | 575 | 6 |
| | 600 | 5 |
| | 625 | 5 |
| | 650 | 5 |
| | 675 | 6 |
| | 700 | 8 |
| dark | | 6 |

## 230706 - 575nm

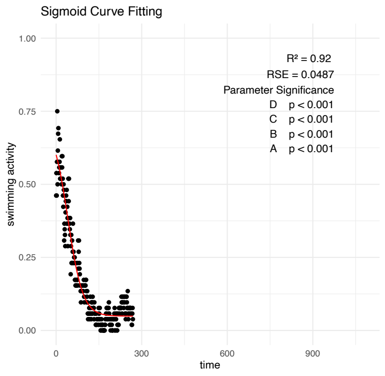


## 230523 - 575nm

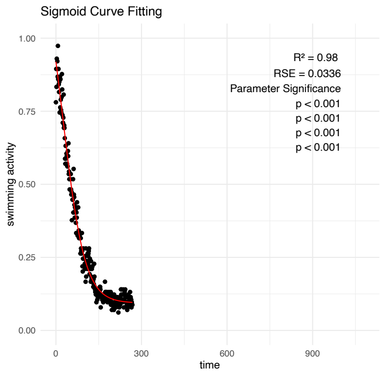


## 230515 - 575nm

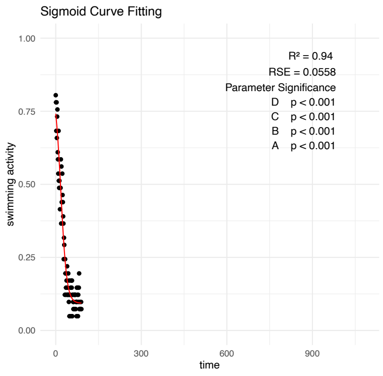


## 230807 - 600 nm

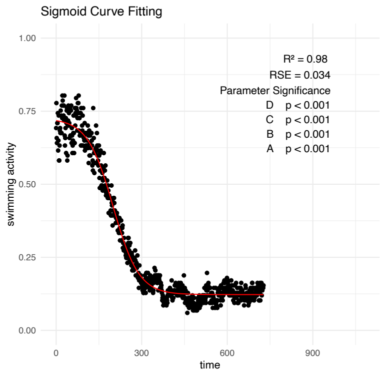


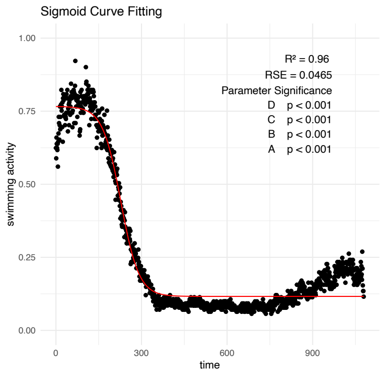


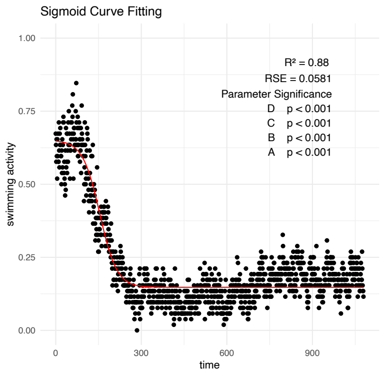


## 230710 - 625 nm

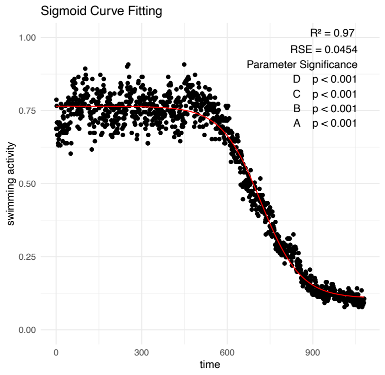


## 230913 - 625 nm

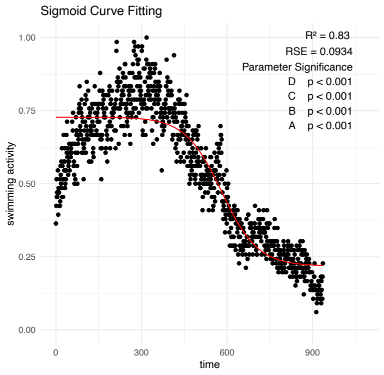


## 230807 - 625 nm

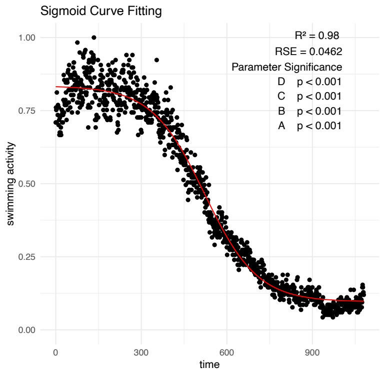


### 230817 - 650 nm **no fit**

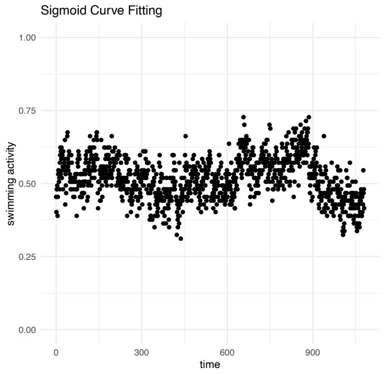


## 230807 - 650 nm

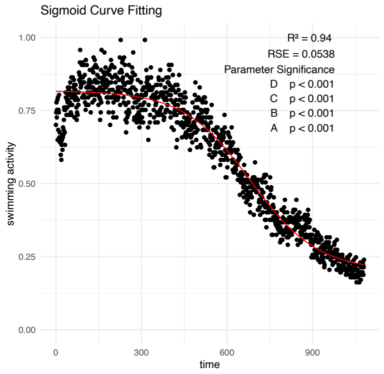


## 230629 - 650 nm

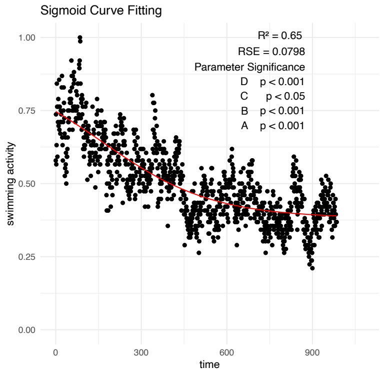

### Exp1 675 nm

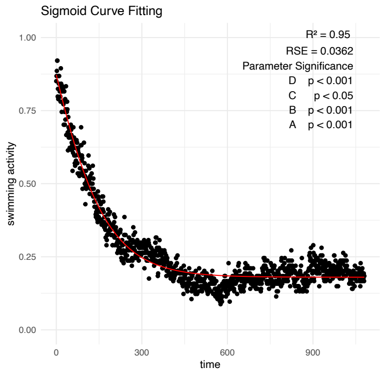


### Exp6 675 nm, **no fit**

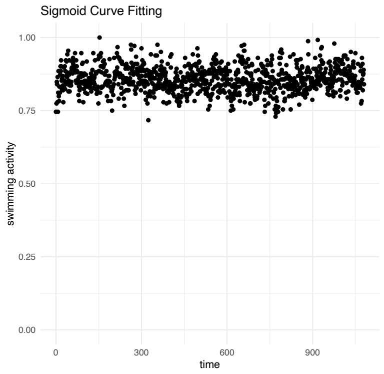


### Exp5 675 nm

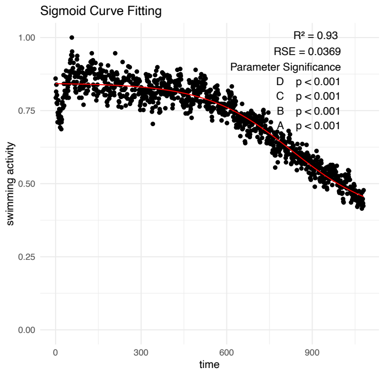


### Exp5 - 700 nm **no fit**

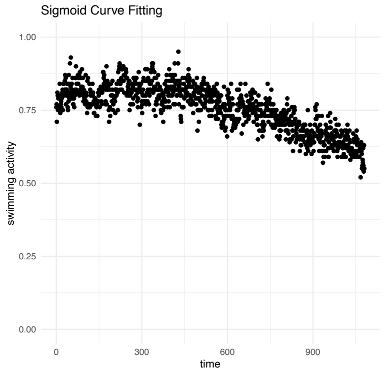


### Exp6 - 700 nm

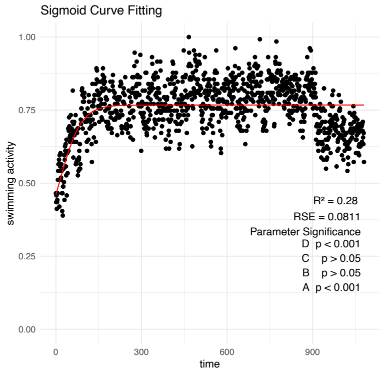


## 230817 - 700 nm

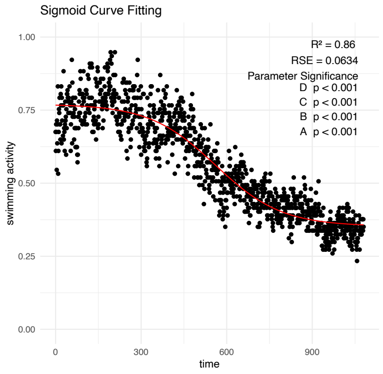


### Exp3 - dark

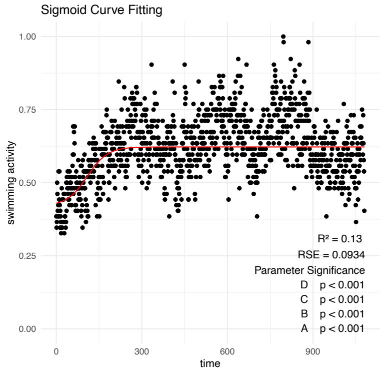


### Exp5 - dark

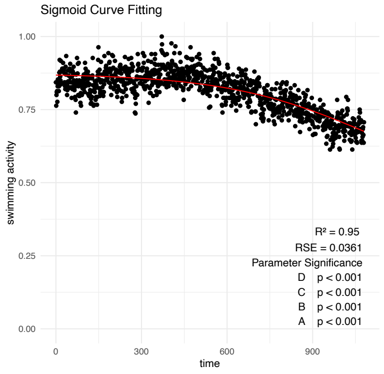


### Exp7 - dark

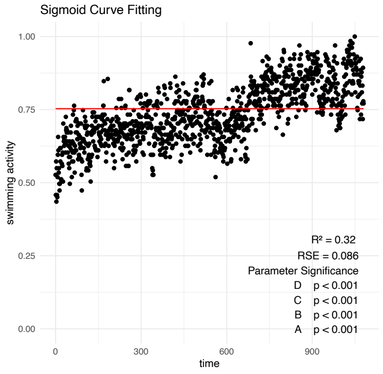
